# Ultrastructure of viruliferous *Javesella pellucida* transmitting Festuca leaf streak virus (genus *Cytorhabdovirus*)

**DOI:** 10.1101/258483

**Authors:** Thorben Lundsgaard

## Abstract

The ultrastructure of cells in the head and thorax from viruliferous *Javesella pellucida* transmitting Festuca leaf streak virus was studied. Aggregates of nonenveloped nucleocapsid particles were observed at the periphery of viroplasms located in cytoplasm of salivary gland cells, fat cells, and nerve cell bodies. Aggregates of nucleocapsid particles, not associated with viroplasms, were seen within a distance of about 1 μm from the basal lamina of salivary glands. Enveloped virions, singly or aggregated, were observed in nerve cell axons and/or dendrites.

## Introduction

Festuca leaf streak virus (FLSV) belongs to genus *Cytorhabdovirus* in the virus family *Rhabdoviridae* (Walker et al., 2000). In its plant host *Festuca gigantea,* aggregates of enveloped virions accumulate in cisternae of endoplasmic reticulum (ER) near viroplasms (Lundsgaard and Albrechtsen. 1979). FLSV N-protein is present in the viroplasms (Lundsgaard, 1992) and G-protein can be detected in the ER (Lundsgaard, 1995) of infected cells, thus supporting the concept that viroplasms are the site for viral nucleocapsid synthesis with subsequent virion maturation at the surrounding ER membranes.

FLSV is transmitted by the delphacid planthopper *Javesella pellucida* (F.) in a persistent manner (Lundsgaard, 1999). Two other viruses in the genus *Cytorhabdovirus* are transmitted in the persistent manner by delphacid planthoppers, namely Barley yellow striate mosaic virus (bYSMV) (Conti, 1980) and Northern cereal mosaic virus (NCMV) (Yamada and Shikata, 1969). Serological relationship has been shown between BYSMV and NCMV (Milne et al., 1986), but not between FLSV and BYSMV or NCMV (Lundsgaard, 1984). Aggregates of enveloped BYSMV virions accumulate in membranous sacs close to viroplasms in the cytoplasm of salivary glands from infected *Laodelphax striatellus* (Fallén). Thinner rods, believed to be nucleocapsids, are also present in salivary glands (Conti and Plumb, 1977). In the same planthopper species, but infected with NCMV, non-enveloped nucleocapsids accumulate near viroplams in salivary gland cells (Shikata, 1979) and in fat cells (Shikata and Lu, 1967). As nothing is known about the ultrastructure of FLSV in its planthopper vector, the present study, dealing with electron microscopic examination of cells in the head and thorax from viruliferous *Javesella pellucida,* was initiated and presented here.

## Materials and methods

Specimens of *Javesella pellucida* (Fabricius) were collected in a woodland locality from *Festuca gigantea* (L.) Vill. plants infected with FLSV. The planthoppers were propagated in greenhouse on FLSV-infected *F. gigantea.* Nymphs collected from infected host plants were caged individually on a healthy barley seedling for 4 days. To select the viruliferous planthoppers, they were transferred individually to a closed, transparent plastic tube (5 ml capacity) containing 0.5 ml water and a 5 cm leaf segment of *Poa trivialis* and kept in a greenhouse for 10 days at 10-15 °C. During this period, the barley test plants were incubated in greenhouse at 20-25 °C. At the end of the period, the barley test plants with symptoms indicated which of the planthoppers were viruliferous.

For fixation, dehydration, and embedding, the method of Berryman & Rodewald (1990) was essentially followed. All incubation steps were done at 5-10 °C. The viruliferous planthoppers were immersed in fixative (3% formaldehyde, 0.3% glutaraldehyde, 100 mM phosphate buffer, pH 7.0), thus head was severed and kept in fixative for at least 2 h. After washing (3.5% sucrose, 0.5 mM CaCl_2_, 100 mM phosphate buffer, pH 7.4), the aldehyde groups were quenched with 50 mM NH_4_O in the washing solution for 1 h. The specimens were then washed (3.5% sucrose, 100 mM maleate buffer, pH 6.5) before post-fixation with 2% uranyl acetate in the same maleate buffer for 2 h. After dehydration in acetone, the specimens were infiltrated in LR Gold (The London Resin Co., England) containing 0.5% benzoin methyl ether (3 changes, 1 day each) and polymerised under near ultraviolet light at room temperature. Control specimens from planthoppers not exposed to FLSV were prepared in the same way. Thin sections (60-70 nm) were collected on 200-mesh nickel grids, stained with saturated uranyl acetate for 5 min and lead citrate (1 mM, pH 12) for 1 min, and then examined in a JEOL 100SX transmission electron microscope operated at 60 kV.

## Results

Head and thorax of *J. pellucida* is mainly occupied by four types of tissue: fat cells, salivary gland cells, brain tissue, and muscle tissue. Virus induced structures were observed in all except muscle tissue. Granular masses with diameters up to 7 μm were present in cytoplasm of fat cells, salivary gland cells, and nerve cell bodies of viruliferous planthoppers (Figure 1). Aggregates of non-enveloped rod-shaped particles, 200-220 nm long and 25-30 nm in diameter, were usually observed at the periphery of these masses (Figure 1, insert). The granular masses are believed to be viroplasms, the sites for viral replication of nucleocapsids.

**Fig. 1.**
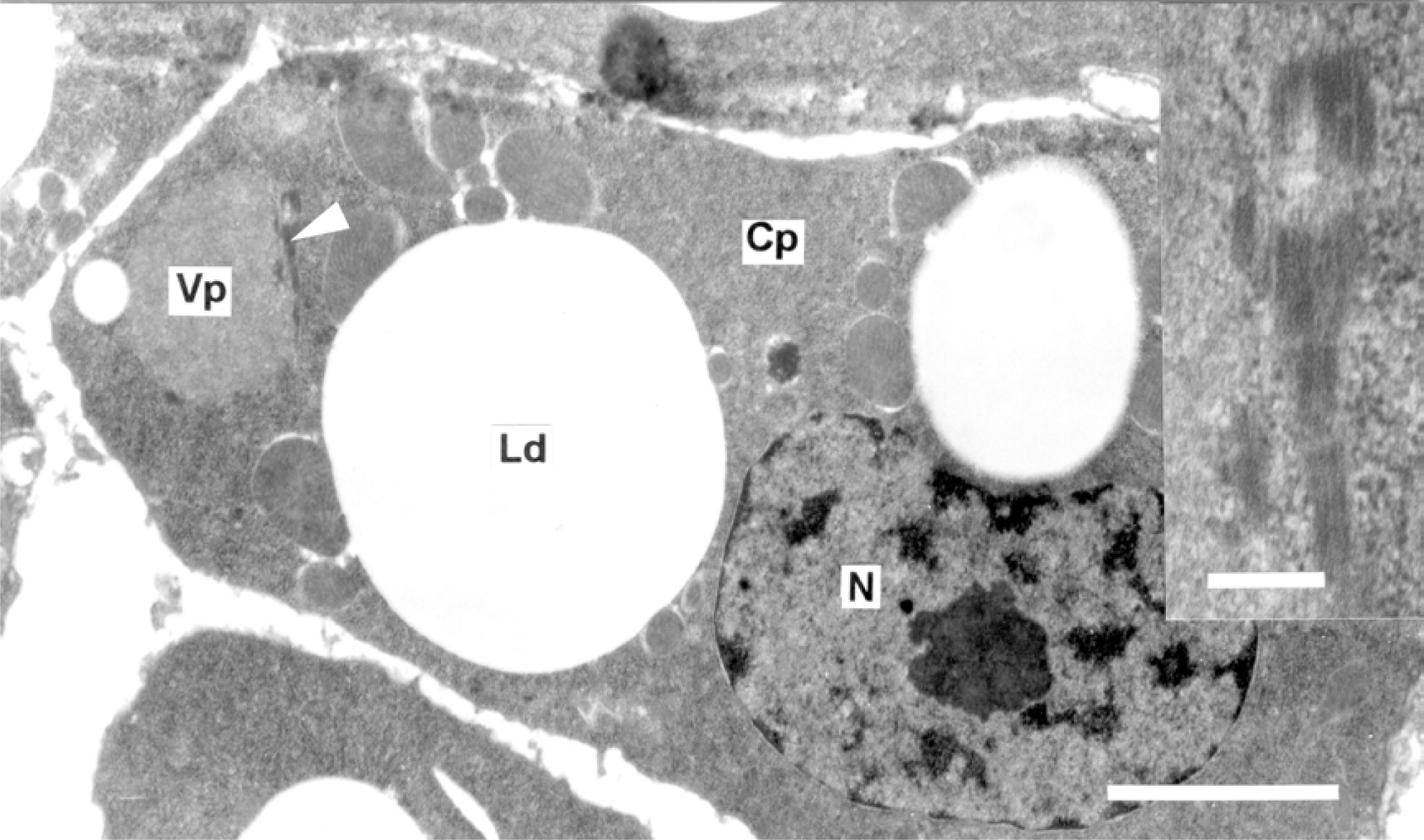
Section through a fat cell from a viruliferous *Javesella pellucida* transmitting Festuca leaf streak virus. Nucleocapsid particles cut nearly longitudinally (arrowhead) are seen at the periphery of a viroplasmic area **(Vp)** in the cytoplasm **(Cp**). The insert in the upper right hand corner is an enlarged picture of the nucleocpasids marked with arrowhead. **Ld** = lipid droplet, **N** = nucleus. Bar marker represents 2 μm in main picture and 200 nm in the insert.

Aggregates of nucleocapsids not associated with viroplasm were often observed within a distance of 1 μm from the basal lamina of salivary gland cells (Figure 2). Typically, these aggregates are separated from the cytoplasm by a membrane, which looks like (and sometimes seen continuous with) the plasma membrane. The cytoplasm in these nucleocapsid-containing sacs has low ribosome content and does not contain endoplasmic reticulum. Probably, these nucleocapsid aggregates are in a translocation process in or out from the cell in concern.

**Fig. 2.**
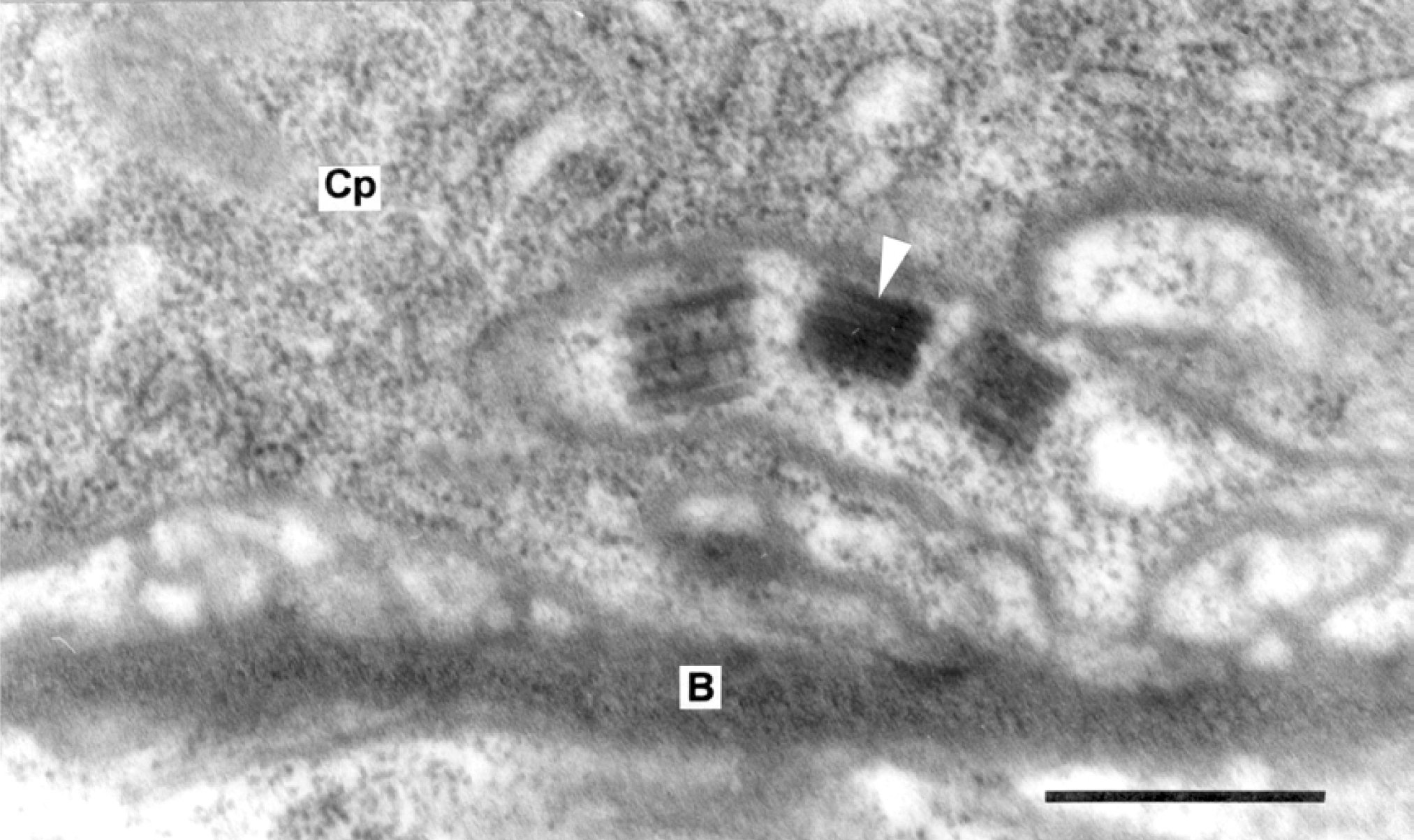
Section through a salivary gland cell from a viruliferous *Javesella pellucida* transmitting Festuca leaf streak virus. Aggregates of nucleocapsid particles cut longitudinally (arrowhead) are seen in the cytoplasm **(Cp)** close to the plasma membrane and basal lamina **(B).** Note the membrane (probably plasma membrane infolding) enclosing the aggregates. Bar marker represents 500 nm.

Most of the brain tissue consists of nerve cell axons and dendrites characteristic by presence of mitochondria and microtubules, only. It was not possible in the present study to distinguish between axons and dendrites. Aggregates of enveloped FLSV virions, 220 nm long and 45-50 nm in diameter, were observed in axons/dendrites from viruliferous insects (Figure 3). Sometimes, a membrane could be seen enclosing such virion aggregates as also seen in Figure 3. When cut in transverse (Figure 3, insert), an electron-dense inner nucleocapsid core can be seen surrounded by a membrane (electron translucent layer) in which the glycoprotein projections (the outermost electron-dense layer) are inserted. As described above, viroplasms are present in some nerve cell bodies, so, nucleocapsids synthesised here are probably enveloped at some nerve cell membrane (not observed) and then translocated as enveloped virions through axons/dendrites to other parts of the insect.

**Fig. 3.**
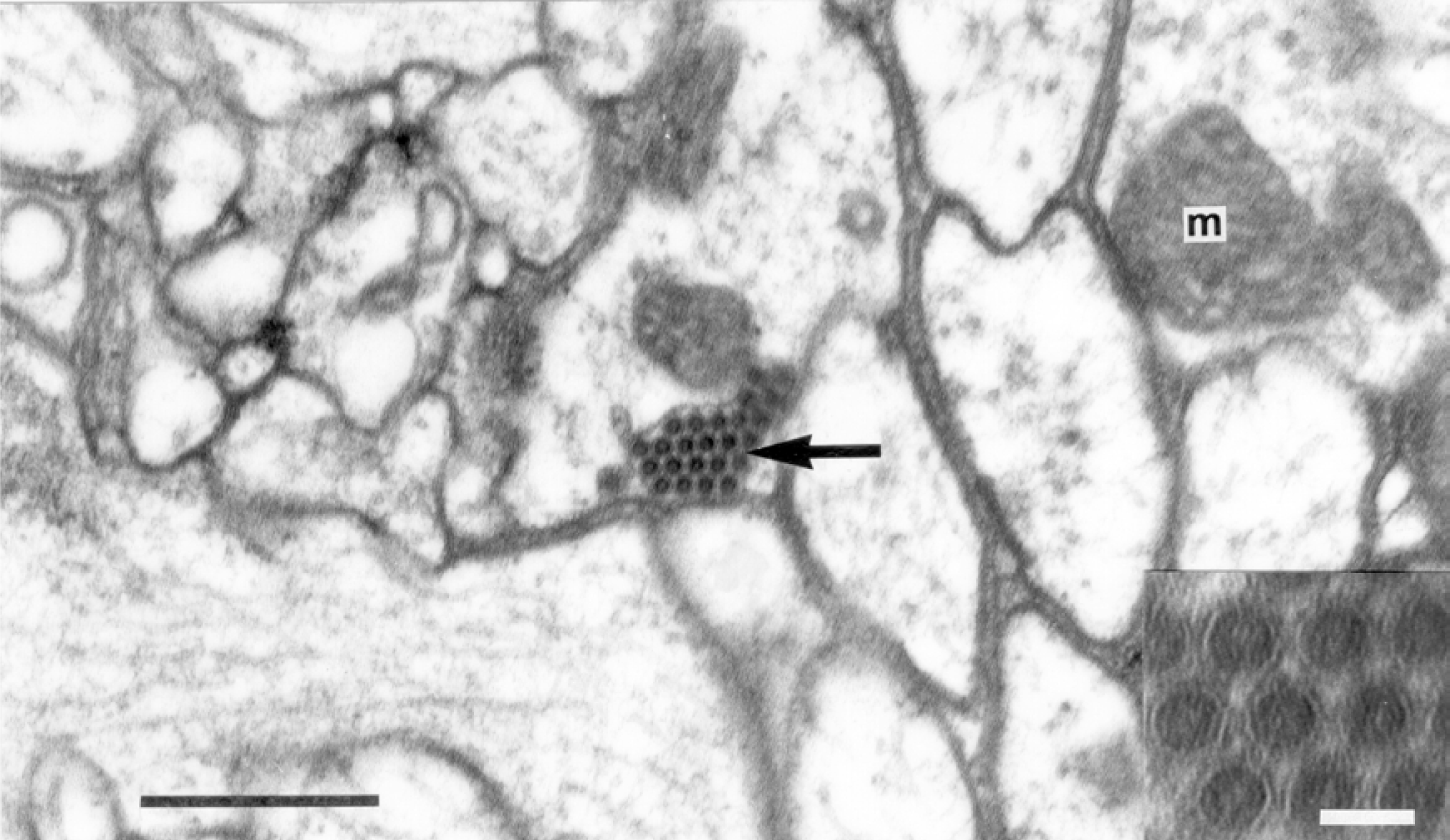
Section through brain tissue from a viruliferous *Javesella pellucida* transmitting Festuca leaf streak virus. An aggregate of enveloped FLSV particles (arrow) is seen enclosed by a membrane in the axonic or dendritic part of a nerve cell. The insert in the lower right hand corner shows an enlarged picture of virus particles cut transversely. The electron-dense inner nucleocapsid core is surrounded by a membrane (electron translucent layer) in which the glycoprotein projections (the outermost electrondense layer) are inserted. **m** = mitochondrion. Bar marker represents 500 nm in main picture and 50 nm in the insert.

## Discussion

The dimensions of FLSV virions in brain tissue reported here are smaller than reported for FLSV virions in plant host cells (Lundsgaard and Albrechtsen, 1979). This difference can be explained by use of methacrylate resin (LR Gold) in the present study compared to use of epoxy resin (Epon) in the former study. Epoxy resins does not shrink during polymerisation, but methacrylate resins shrink up to 20% during polymerisation (Glauert, 1975).

The ultrastructure of plant cells (Lundsgaard and Albrechtsen, 1979) and planthopper cells (present study) infected with FLSV shows that replication takes place in cytoplasmic viroplasms in both host types. In the plant host, nucleoprotein just synthesised is enveloped at the surrounding G-protein-containing endoplasmic reticulum (Lundsgaard, 1995), but in salivary gland and fat cells of the vector host, the surrounding ER is not competent for enveloping FLSV. In a preliminary investigation of infected salivary gland cells for presence of G-protein antigen using the same procedure as used for plant cells (Lundsgaard, 1995), no G-protein could be detected in these cells (unpublished).

The surface G-protein of rhabdoviruses is believed to play an important role for host recognition. The uptake of FLSV in healthy planthoppers is presumed to take place in the alimentary tract with following recognition of G-protein at some cell surface receptors. So, it makes sense that the virus source (plant tissue) contains enveloped particles. The vector transmission of virus into healthy plants is believed to take place by delivering saliva with virus directly into the plant sieve tubes, which are part of the plant symplast. In such a scenario there is no exposure of virus to cell surfaces and thus no need for viral G-protein recognition. For cytorhabdoviruses, the nucleocapsid entity is infectious (Dietzgen, 1995) and nucleocapsids delivered from salivary glands of viruliferous planthoppers have thus the potential for establishing a new infection in the plant host. So, it makes sense that G-protein for enveloping is not produced in salivary glands.

The mechanism of release of virus particles into salivary ducts is not known and the present study did not unveil details for this process. An obvious candidate for delivering the virus into the salivary ducts is the secretory pathway. Careful examination of such secretory vesicles from viruliferous planthoppers in the present study did not show nucleocapsid particles inside, but presence of uncoiled viral nucleoprotein can not be excluded.

The ultrastructure of salivary gland cells from planthoppers infected with the two cytorhabdoviruses BYSMV and NCMV shows that BYSMV particles accumulate both as nucleocapsids as well as enveloped particles, whereas NCMV particles accumulate as nucleocapsids, only. In this respect, FLSV resembles NCMV more than BYSMV. Enveloped FLSV virions were observed in axons and/or dendrites of viruliferous insects in the present study and this is the first time cytorhabdovirus particles have been detected in planthopper brain tissue. The systemic spread of persistent viruses in vectors is believed to take place by hemolymph circulation in the hemocoel (Ammar, 1994). The present study suggests that axons and/or dendrites also could play a role for systemic spread of rhabdoviruses in their insect vectors. Rabies virus in the genus *Lyssavirus* of *Rhabdoviridae* is known to use axonal transport along microtubules to infect the central nervous system of its vertebrate host (Almenar-Queralt and Goldstein, 2001).

Neither viroplasms nor virus particles were observed in cells of *J. pellucida* not exposed to FLSV infection.

